# The *PPE25 (Rv1787) - PE18 (Rv1788) - PPE26 (Rv1789)* gene cluster encodes an interacting protein pair and is involved in immune evasion by *Mycobacterium tuberculosis*

**DOI:** 10.1101/2023.04.07.535983

**Authors:** Ravi Prasad Mukku, Kokavalla Poornima, Korak Chakraborty, Tirumalai R. Raghunand

## Abstract

*Mycobacterium tuberculosis (M. tb)* the causative agent of human tuberculosis, encodes multiple virulence factors to subvert host immune responses. One such class of proteins is encoded by the multigenic *PE_PPE* family which accounts for 10% of its coding potential. A number of these genes occur in clusters, of which the *PPE25(Rv1787)-PE18(Rv1788)-PPE26(Rv1789)* locus alone, is organised in a *PPE-PE-PPE* arrangement. We establish here that this cluster is co-operonic in *M. tb*, and identify for the first time, PPE25::PPE26 as the sole interacting protein pair encoded by this cluster. Recombinant *M. smegmatis* strains expressing *PPE25, PE18*, and *PPE26*, exhibited enhanced survival in THP-1 macrophages, with infected cells displaying increased levels of the anti-inflammatory cytokine *IL-10*, and reduced levels of the pro-inflammatory cytokine *IL-12*. Macrophages infected with recombinant *M. smegmatis* expressing *PPE26* showed increased phosphorylation of the MAP kinase p38, consistent with the known TLR2 binding activity of PPE26. In contrast to strains expressing the individual cluster genes, the recombinant expressing the entire *PPE25-PE18-PPE26* operon showed no change in intra-macrophage CFUs, suggestive of an inhibitory role for the PPE25::PPE26 complex in CFU enhancement. Taken together, our findings implicate the *PPE25-PE18-PPE26* cluster in playing an immune evasion role in the pathophysiology of *M. tb*.

## Introduction

*Mycobacterium tuberculosis* (*M. tb*), the causative agent of human tuberculosis, possesses the ability to thrive and persist within macrophages by altering their immediate antimicrobial actions and evading host immunity (1). Mycobacteria have highly hydrophobic and impenetrable cell walls, and therefore need very specific and efficient mechanisms to secrete the effectors needed for their immune evasion function. Their genomes encode an atypical type VII secretion apparatus, commonly known as the ESX system, and also contain transporters known in other bacteria (2). A growing body of evidence has revealed that the multigenic PE_PPE proteins, which comprise close to 10% of the coding potential of the *M. tb* genome, are either secreted from or are associated with the mycobacterial cell envelope, and are linked to antigenic diversity, immune evasion, and mediating host-pathogen interactions (3). Our understanding of the molecular basis of their functioning however, remains limited.

Interestingly, the 12 gene ESX-5 locus contains two clusters of members of the PE_PPE family of proteins, including a three gene segment encoding PPE25 (*Rv1787*), PE18 *(Rv1788)*, and PPE26 (*Rv1789*) (4, 5). The *M. tb* Δ*PPE25-PE19* mutant containing a deletion in 5 PE_PPE genes encoded in the *esx5* locus was shown to be avirulent in immunocompetent mice. This strain however remained strongly immunogenic and showed a greater protective efficacy against *M. tb* infection than BCG (6). This 5 gene deletion strain also showed reduced virulence in murine macrophages, and in a SCID mouse model of *M. tb* infection (5). Several bacterial processes, including replication, vacuole acidification, and phagolysosome fusion in macrophages, were affected when the gene homologous to *PPE25* was deleted from *Mycobacterium avium* (7). PPE25 has been used as a strong T-cell epitope for a DNA vaccine against *M. tb* (8). Moreover, the expression of the *PPE25, PE18* and *PPE26* was observed to be upregulated under conditions of hypoxia/ dormancy, and NO stress (9, 10). Also, PPE25 and PPE26 were shown to influence cytokine secretion and increase the survival of *M. smegmatis* in mouse macrophages (11). Taken together, these observations are strongly suggestive of a significant role for this gene segment in the pathophysiology of *M. tb*, leading us to functionally evaluate this cluster for its role in *M. tb* pathogenesis.

## Materials and methods

### Bacterial strains, media and growth conditions

*M. smegmatis* mc^2^6 and *M. tb* H37Rv were cultivated in Middlebrook, 7H9 broth and 7H10 agar (Difco) containing albumin dextrose complex (5 g BSA, 2 g glucose, and 0.85 g NaCl/L), 0.5% (v/v) glycerol, and 0.5% (v/v) Tween 80. Luria Bertani medium was used to cultivate *E. coli* strains - both *E. coli* and mycobacteria were cultured by shaking at 37° C. When required, 200 µg/ml of ampicillin and 50 µg/ml of kanamycin (for *E. coli*) and 15 µg/ml of kanamycin (for mycobacteria) were added to the culture medium.

### Sub-cellular localisation and Proteinase K sensitivity assay

In order to determine the subcellular location of these proteins, the *M. tb PPE25, PE18, PPE26* ORFs were cloned between *BamHI* and *EcoRI* sites of pJEX55 (12) and the recombinant plasmids were electroporated into *M. smegmatis*. Recombinants expressing c-myc tagged PPE25, PE18, PPE26 were harvested at the logarithmic phase of growth, washed and resuspended in PBS. The samples were divided into two identical aliquots and incubated at 37°C for 30 min with or without Proteinase K (Sigma). The reaction was stopped by adding 2 mM EGTA, and sub-cellular fractions of these samples were isolated as described in (13). The individual fractions were separated using denaturing SDS PAGE, and the fusion proteins were detected by western blotting with anti-c-myc antibodies (sc40, Santa Cruz).

### Expression of PPE25, PE18, and PPE26 in M. smegmatis

*M. tb PPE25, PE18, PPE26* ORFs were amplified using gene-specific primers (Table S1) and cloned between the *BamHI* and *EcoRI* sites of pMV261 (14). These constructs were then transformed into *M. smegmatis* for overexpression.

### Expression and purification of PPE25 and PPE26

The full-length *M. tb PPE25* and *PPE26* ORFs were cloned as C-terminal 6xHis-tagged fusion in pET22b, and N-terminal Glutathione-S-Transferase (GST) tagged fusion in pGEX6p1 respectively. Following sequence confirmation, the constructs were transformed into *E. coli* BL21 (DE3). For induction, cultures were allowed to grow to an OD of 0.5, and induced with 0.5 mM IPTG. His-tagged PPE25 was purified using Ni-NTA affinity chromatography and GST tagged PPE26 was purified using glutathione agarose beads. The purified proteins were used for pull-down experiments.

### In vitro growth kinetics

To functionally characterise *M. tb PPE25, PE18*, PPE26, their ORFs were amplified from *M. tb* H37Rv genomic DNA using gene specific primers (Table S1), cloned between the *BamHI* and *EcoRI* sites of pMV261 and transformed into *M*.*smegmatis*. To examine their growth patterns, recombinant *M. smegmatis* strains were grown until late exponential phase, diluted to an OD of 0.2, and cultured in Middlebrook 7H9 broth containing 15 µg/ml kanamycin. Growth curves were generated by CFU measurements and plotted against time. At each designated time point, cultures were harvested for RNA extraction and gene expression analyses performed. All CFU enumeration assays were performed in the presence of 15 μg/ml kanamycin.

### Macrophage infection

THP-1 culture and all infections, were performed as described in (15).

### Real-time RT-PCR analysis

The expression profiles of *PPE25, PE18 PPE26* in *M. tb* and in recombinant strains of *M. smegmatis* expressing pMV261-*PPE25*, pMV261-*PE18* pMV261-*PPE26* as a function of growth, were determined using real-time transcript measurements as described in (15), using *PPE25, PE18, PPE26* specific primers (Table S1). Gene specific transcript levels in *M. smegmatis* were normalised to the *sigA* transcript in each sample, and in *M. tb* to the 16s rRNA transcript. In *M. smegmatis* the relative fold change in transcript levels at each time point was calculated with respect to the levels at 4 h. In the *M. tb* expression profiles, the relative fold change in transcript at each OD point was calculated with respect to the levels at OD 0.8. Quantitation of host cytokines and *iNOS2* transcript levels was performed as described in (15), using primers specific to *iNOS2, IL-10, IL-12* (Table S1). The levels of each mRNA were normalised to the transcript levels of GAPDH and β-actin. Relative fold change was calculated with reference to macrophages infected with *M. smegmatis* expressing the empty vector.

### Western blotting

The levels of total and phosphorylated p38 in infected THP-1 macrophages were determined by preparing cell lysates in RIPA lysis buffer. SDS-PAGE and western blotting were performed as described in (15). The normalised levels of phosphorylated p38 in control samples (pMV261) were assigned a value of 1, and the fold change in the levels of these proteins in the test samples (PPE25, PE18, PPE26) were derived with respect to the control.

### Co-transcriptional analysis

Total RNA extracted from exponentially growing *M. tb* H37Rv was used to assess the transcriptional state of the *PPE25-PE18-PPE26* cluster. RNA treated with DNAse I was used as a template for cDNA synthesis with the primers *PPE26* R1 (Table S1). The cDNA was utilised for PCR amplification with gene-specific primer combinations. The analysis includes the appropriate negative controls (-RT).

### Co-immunoprecipitation

To perform *in vivo* pull down assays, 150 ml of independent *M. smegmatis* cultures co-transformed with pJEX55+pSCW54, pJEX55-*PPE26+*pSCW54, pJEX55+pSCW54*-PPE25*, pJEX55-*PPE26+*pSCW54*-PPE25* were grown in Middlebrook 7H9 broth containing kanamycin and hygromycin to log phase and induced with 0.2% acetamide for 6 h. Induced cells were pelleted, resuspended in 1.2 ml PBS containing protease inhibitors and lysed by sonication. All further steps were performed as described in (16).

### Mycobacterial Protein Fragment Complementation (MPFC) assay

To assess the interaction of PPE25-PE18, PE18-PPE26 and PPE25-PPE26 *in vivo, PPE25, PE18* and *PPE26* were cloned into the *E. coli* - *Mycobacterium* shuttle vectors pUAB400 and pUAB300 respectively. Both constructs were co-transformed in *M. smegmatis* mc^2^155 and the interaction assay was performed as described (17). Appropriate positive (pUAB100+pUAB200) and negative (pUAB100+pUAB400, pUAB400+ pUAB300) controls were included in the assay.

### In silico analyses

All *M. tb* sequences were obtained from the Mycobrowser database (https://mycobrowser.epfl.ch/). The Dense Alignment Surface (DAS) method (18) and Kyte & Doolittle (19) algorithms were used to identify trans-membrane domains and regions of hydrophobicity respectively.

## Results

### PPE25, PE18 and PPE26 are co-operonic and the PPE25 and PPE26 gene products interact

The *PPE25-PE18-PPE26* cluster, is the only one of 40 clusters of PE/PPE genes in the *M. tb* genome that has a *PE* gene that is directly flanked by two *PPE* genes, in a tandem arrangement (20). Its predicted co-operonic status, is based on the short spacing between the genes, with 78 bp separating *PPE25-PE18*, and 13 bp separating *PE18-PPE26* (Fig. 1A). To confirm this prediction, RT-PCR experiments were carried out with RNA isolated from log phase grown *M. tb* H37Rv using primer pairs (Table S1) intended to amplify ORF and junction-specific areas of the cluster (Fig. 1A). The sizes of the amplicons obtained (*PPE25*-200 bp, J1-200 bp, *PE18*-200 bp, J2-200bp, *PPE26*-200 bp), and the absence of amplification products in the -RT control lanes, establish that this gene cluster is transcribed as a mono-cistronic message (Fig. 1B).

**Figure 1:**
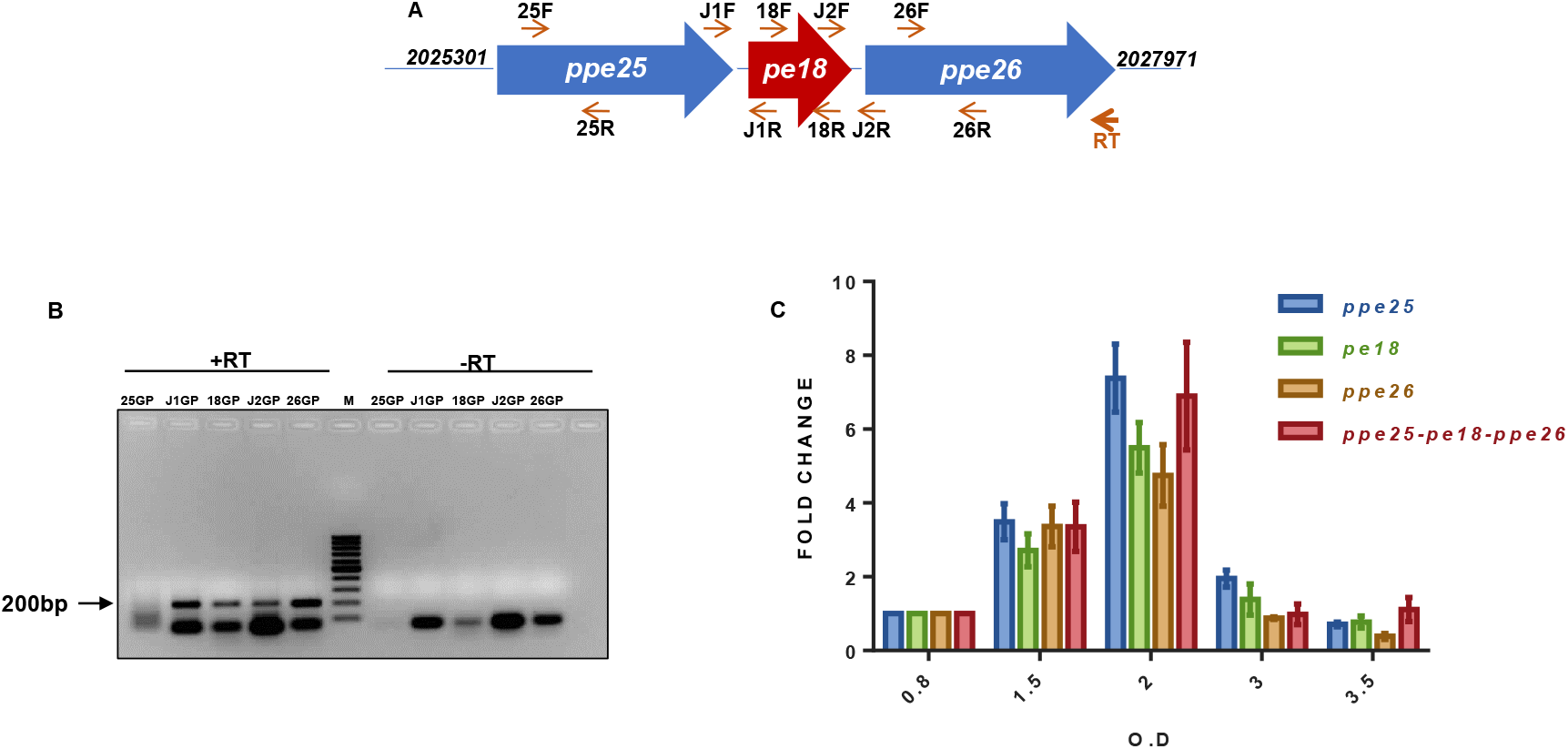
Analyses of operonic organisation of the *PPE25-PE18-PPE26* gene cluster. A) The genomic organisation of the *PPE25-PE18-PPE26* region of the *M. tb* genome is shown schematically to highlight the positions of the gene (G) and junction (J) specific primers used in the experiment. (B) RT-PCR products amplified from the *PPE25-PE18-PPE26* cluster: lane 1: (25F+25R), lane 2: (25JF+18JR), lane 3: (18F+18F), lane 4: (18JF+26JR), lane 5: (26GF+26GR), lane 6: 100 bp DNA ladder (M), lanes 7-11: -RT controls for the respective primer pairs. (C) Real-time RT-PCR quantification of *PPE25, PE18, PPE26* and *PPE25-PE18-PPE26* transcripts as a function of *M. tb* H37Rv growth. Transcript levels are shown relative to the cognate gene’s mRNA levels at 0.8 OD, which is given a value of 1. All findings reported are based on at least two biological replicates.

This observation was further validated by examining the expression levels of the three genes as a function of growth in *M. tb* H37Rv. All the genes were observed to be coordinately expressed (Fig. 1C), confirming the operonic status of this cluster. It is well-known that genes expressed in a single operon have related functions, and their products are most likely to physically interact with each other. To test this possibility, we performed a pairwise screen for interaction between PPE25+PE18, PPE26+PE18, and PPE25+PPE26 using the mycobacterial protein fragment complementation (M-PFC) assay (17). In this screen, the cluster genes were expressed as murine dihydrofolate reductase domain F [1, 2] and F [3] fusions, and then co-transformed into *M. smegmatis* mc^2^155. Co-transformants for the PPE25-PPE26 pair, were observed to be resistant to 20 µg/ml of trimethoprim (Fig. 2A), demonstrating that PPE25 and PPE26 interact *in vivo*. No growth was observed in the negative control samples and co-transformants for the PPE25+PE18 and PPE26+PE18 pairs, implying that this cluster encodes only one pair of interacting proteins, namely PPE25:PPE26. To confirm this interaction, we performed an *in vivo* pull-down in *M. smegmatis*, which lacks this gene pair. PPE25 and PPE26 were shown to co-precipitate in this experiment (Fig. 2B), proving that the two proteins indeed interact *in vivo*; we ensured equal loading for each sample (Figure. S1). This interaction could not be tested *in-vitro* since PPE25 cloned with a C-terminal 6xHis tag could be expressed but not purified (Fig. S2A). We however were able to partially purify an N-terminal GST tagged PPE26 (Fig. S2B).

**Figure 2:**
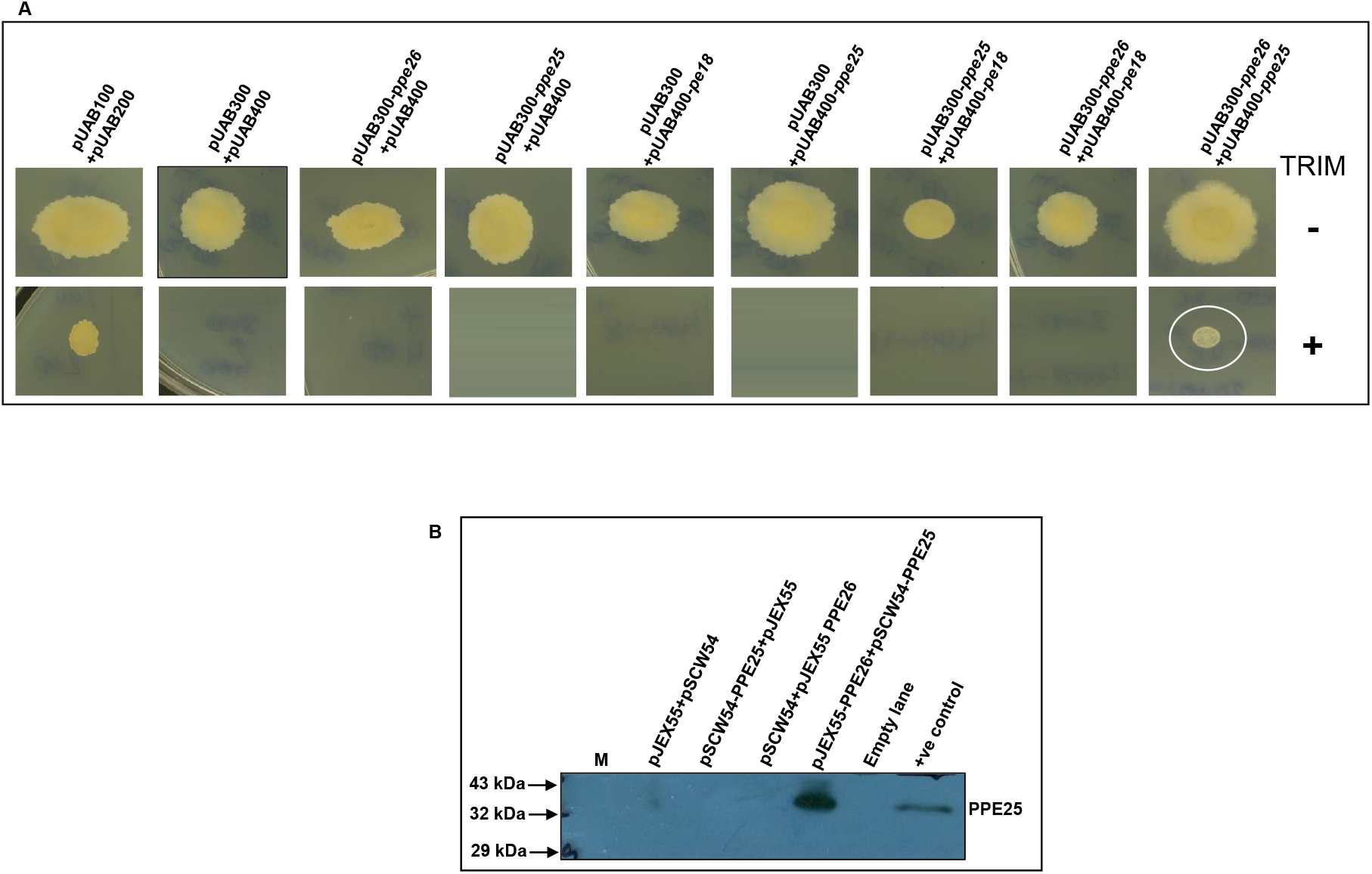
Protein-protein interaction in the *PPE25-PE18-PPE26* gene cluster. (A) M-PFC analysis of pair-wise interaction between PPE25+PPE26, PPE25+PPE18, and PPE26+PPE18. The image depicts the growth pattern of *M. smegmatis* co-transformants on Middlebrook 7H11 agar plates with (+) or without (-) 20µg/ml trimethoprim (TRIM). The circled sample marks positivity for PPE25-PPE26 interaction. (B) Immunoblot of PPE25-PPE26 co-IP reactions from *M. smegmatis* expressing PPE26-c-myc and PPE25-6xHis. An anti-c-myc antibody was used to pull-down the complex, and the blot was probed with an anti-His antibody. All findings reported are based on at least two biological replicates.

### Recombinant strains of M. smegmatis expressing PPE25, PE18, and PPE26, show a survival advantage in macrophages

To investigate the functions of PPE25, PE18, and PPE26 in mycobacterial pathophysiology, we used *M. smegmatis* as a surrogate model, the genome of which lacks homologues for the vast majority of PE-PPE proteins, including PPE25, PE18, and PPE26. First, recombinant *M. smegmatis* strains expressing c-myc tagged PPE25, PE18, PPE26 were subjected to Proteinase K treatment and sub-cellular fractionation to determine the subcellular location of each protein. Each fraction’s loading was ensured to be equal (Fig. S3 A, B, C) before their spatial distribution was analysed. Western blotting showed that all three proteins were localised to the cell envelope, and their signals were greatly reduced in the cell wall-associated fractions in the Proteinase K treated samples as compared to those that were untreated (Fig. S3 D, E, F). This showed that PPE25, PE18, and PPE26 were exposed to the cell surface, allowing the use of this model to evaluate their function as modulators of pathogenicity *via* their interactions with the host. Their cell-envelope localisation was consistent with the computationally predicted presence of transmembrane and hydrophobic stretches in these proteins (Fig. S4A, B). Next, we investigated the possible functions of the proteins in modulating host-pathogen interactions using constitutively expressed *PPE25, PE18*, and *PPE26* in *M. smegmatis*. Over-expression of these genes did not cause growth defects *in vitro*, as shown by growth profiling of *M. smegmatis* strains expressing *PPE25, PE18, PPE26*, the entire operon (*PPE25-PE18-PPE26*), and the empty vector (Fig. S5A, B). Real-time RT-PCR analysis of recombinant *M. smegmatis* strains using gene-specific primers (Table S1) revealed that transcript levels for the three genes were similar, indicating that their expression did not change with growth (Fig. S5C, D, E, F). When we infected THP-1 macrophages with these recombinant strains, we observed that the strains expressing *PPE25, PPE18*, and *PPE26*, showed a survival advantage over the empty vector containing control strain (Fig. 3A). This elevation in CFU counts, was consistent with the reduction in transcript levels of *iNOS2*, which is primarily responsible for control of bacillary load in this model (Fig. 3C). In contrast to this, the strain expressing the full operon, did not show a change in CFU counts compared to the empty vector control (Fig. 3A), which correlated with the absence of change in levels of the *iNOS2* transcript in these cells, (Fig. 3C).

**Figure 3:**
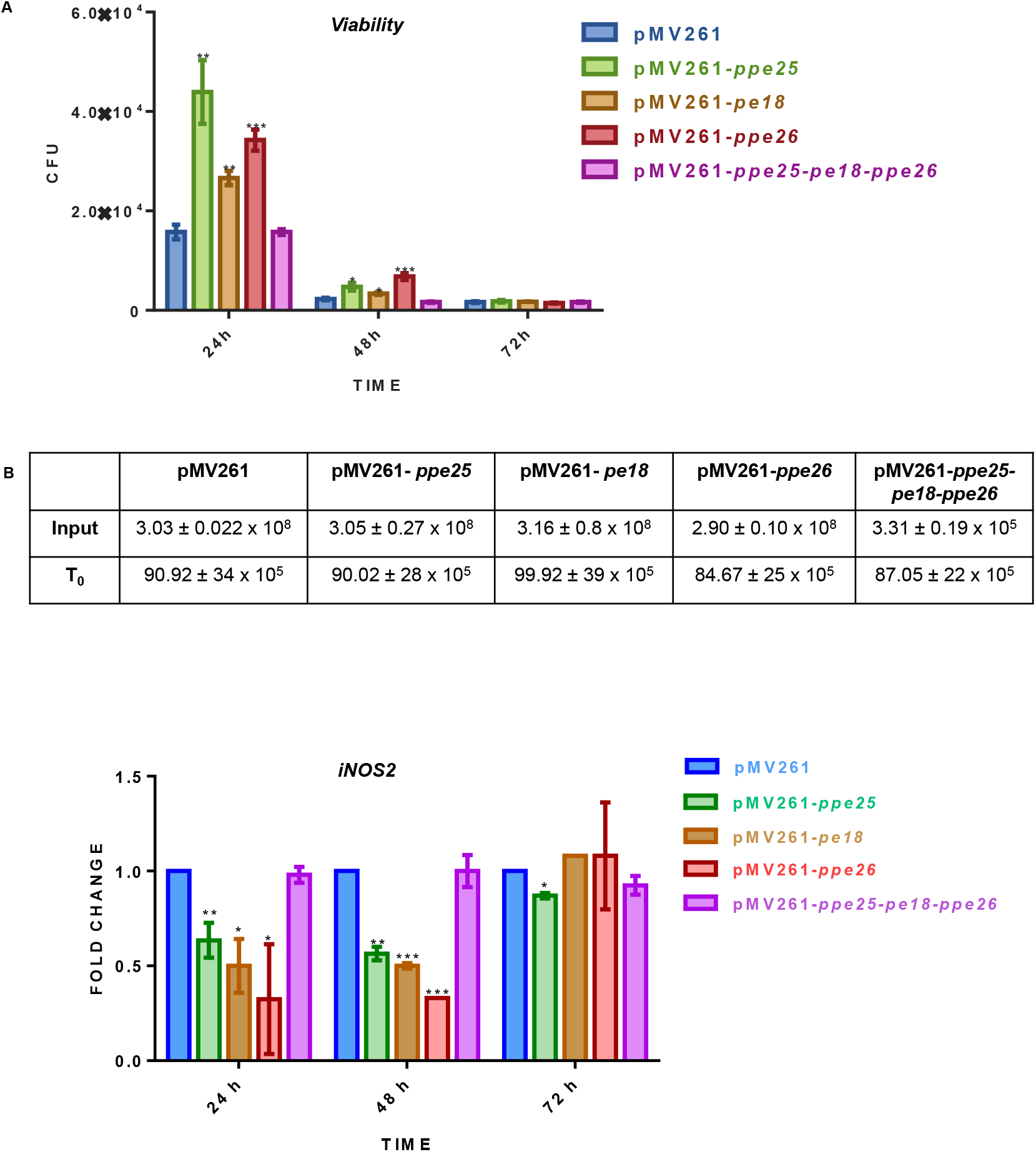
Phenotypes of recombinant *M. smegmatis* expressing*PPE25, PE18*, PPE26 and *PPE25-PE18-PPE26*, in THP-1 macrophages. (A) Post-infection CFU counts of recombinant strains of *M. smegmatis* expressing pMV261, *PPE25, PE18, PPE26* and *PPE25-PE18-PPE26* in THP-1 macrophages at 24 h, 48 h and 72 h. input and post-infection (T_0_) CFU counts of infecting bacilli (±SEM). (C) Real time RT-PCR quantitated *iNOS2* transcript levels in THP-1 macrophages infected with *M. smegmatis* expressing pMV261, *PPE25, PE18, PPE26* and *PPE25-PE18-PPE26*, 24 h, 48 h, and 72 h post-infection. The data is representative of two biological replicates. Error bars represent ± SEM. *p≤0.05, **p≤0.005, ***p ≤0.001. Error bars represent the mean ± SEM of two biological replicates.

### PPE25 and PPE26 modulate the macrophage innate immune response

The levels of numerous cytokines involved in the control of *M. tb* infection (21), were assessed to evaluate the immunomodulatory potential of PPE25, PE18, and PPE26. Among the anti-inflammatory and pro-inflammatory cytokines we measured, IL-10 and IL-12 showed changes in their expression patterns in THP-1 macrophages infected with the recombinant *M. smegmatis* strains. Macrophages infected with recombinants expressing *PPE25, PPE26*, and the entire operon, showed a significant upregulation in transcript levels of the anti-inflammatory cytokine IL-10, but the same was not observed in cells infected with the *PE18* expressing strain (Fig. 4A). Also, the transcript levels of IL-12, a critical pro-inflammatory cytokine, was seen to be significantly down-regulated in macrophages infected with all the recombinant strains (Fig. 4B). Since the interaction of PPE26 with macrophage TLR2 had been described (22), we proceeded to verify if signalling through this receptor was being activated by assessing the phosphorylation status of p38, a marker for TLR2 activation, in the lysates of THP-1 macrophages infected with recombinant *M. smegmatis* expressing *PPE25, PPE18, PPE26*, and the entire operon. An induction in p38 phosphorylation was observed in macrophages infected with *M. smegmatis* recombinants expressing PPE26, and the entire operon, but not in cells infected with recombinants expressing *PPE18* or *PPE25* alone (Fig. 5A, B). This finding validated the observations of Su *et al*. (22), and also opened up a possible mechanistic avenue for the above observations.

**Figure 4:**
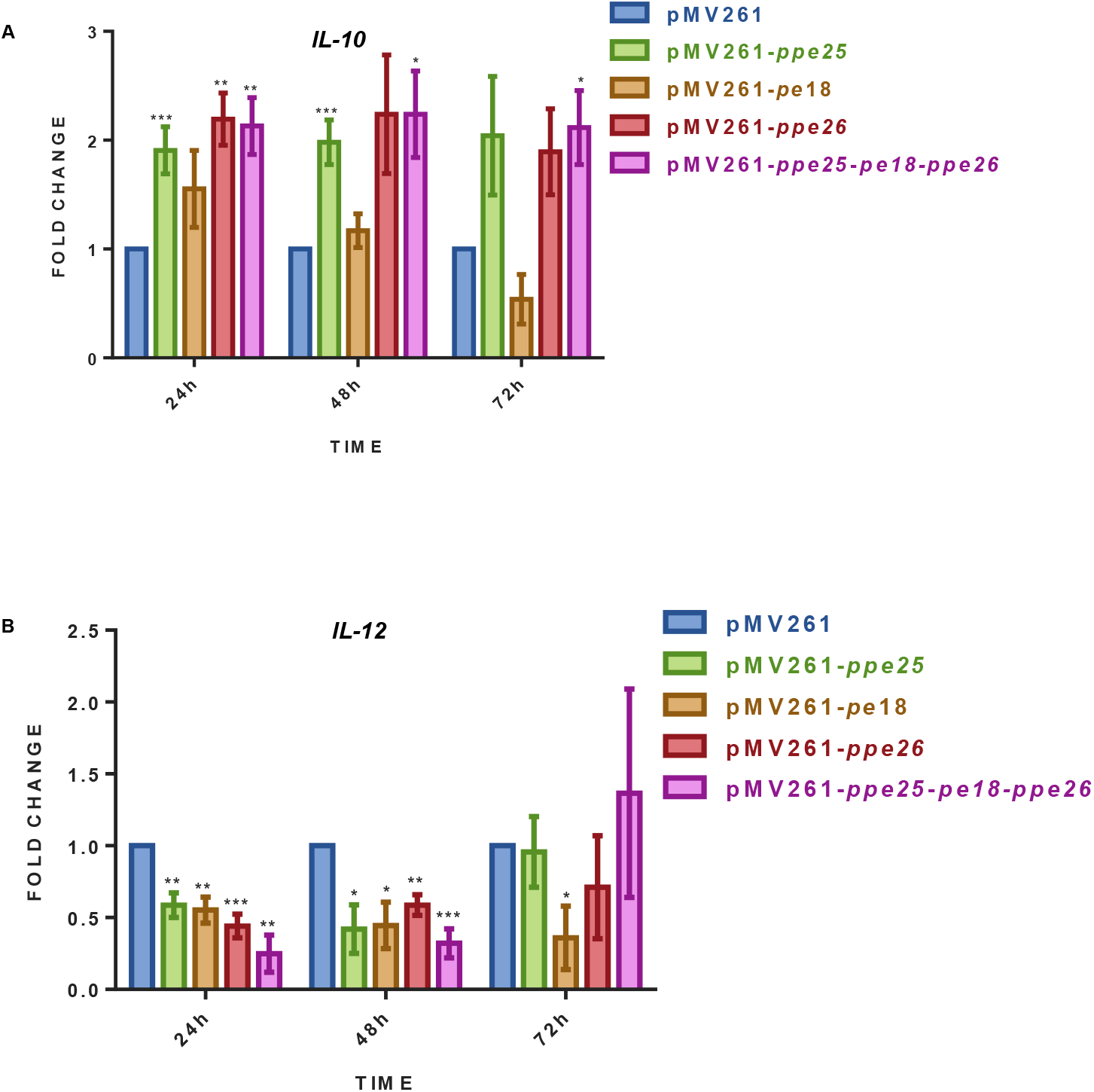
Assessment of immunomodulatory potential. (A) Changes in transcript levels of *IL-10* in THP-1 macrophages infected with recombinant *M. smegmatis* strains expressing *PPE25, PE18, PPE26* and *PPE25-PE18-PPE26*, 24 h, 48 h, and 72 h post-infection. (B) Changes in expression levels of the *IL-12* transcript in the same cells. Error bars represent ± SEM from two biological replicates. *p≤0.05, **p≤0.005, ***p ≤0.001

**Figure 5:**
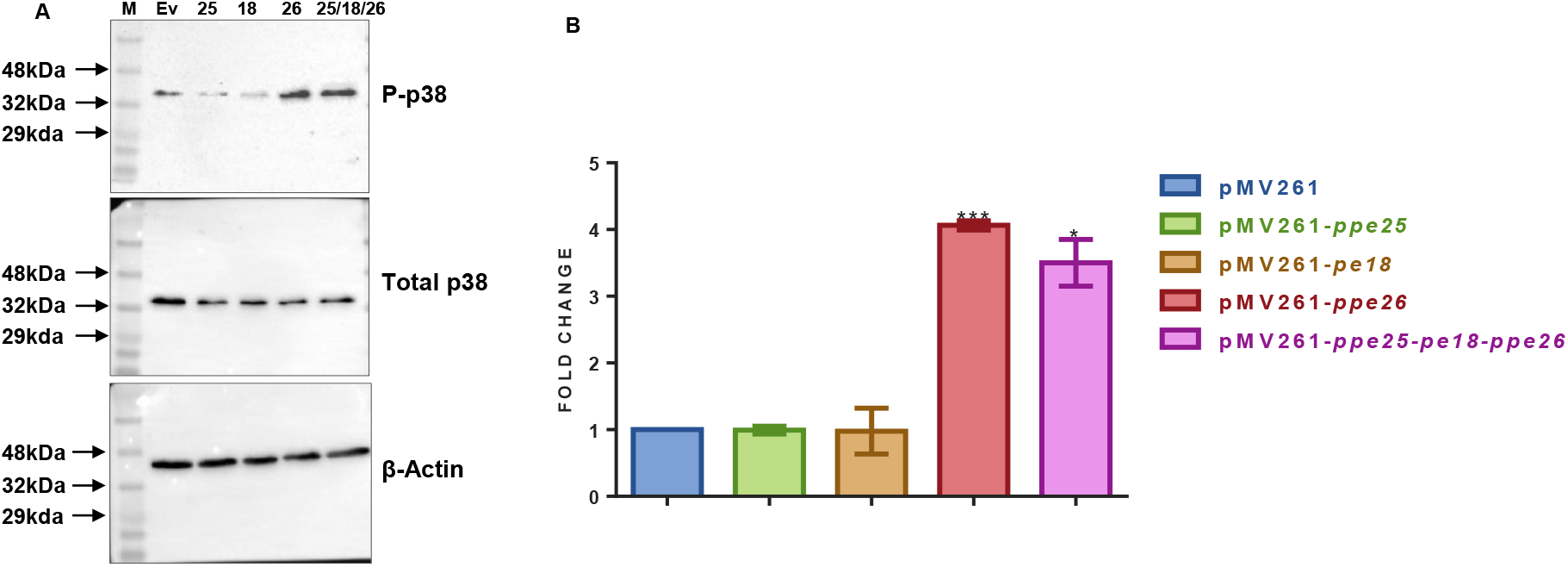
Induction of TLR2 signalling. (A) Western blots depicting levels of total and Phospho-P38, and β-actin in lysates from THP-1 macrophages infected with recombinant strains of *M. smegmatis* expressing pMV261 (Ev), *PPE25* (25), *PE18* (18), *PPE26* (26), and the entire operon (25/18/26). The densitometric quantitation of this data is represented in (B).

## Discussion

In order to survive in the face of a robust host immune responses, pathogenic mycobacteria have developed a wide variety of adaptive strategies (15, 23). This study set out to investigate the involvement of the *PPE25 (Rv1787) - PE18 (Rv1788) - PPE26 (Rv1789)* gene cluster in ESX-5-associated *M. tb* pathogenicity, by assessing its functional characteristics. In addition to their short intergenic spacing, the strong association between the expression patterns of the cluster genes in a dataset exhibiting global transcriptional responses of *M. tb* to inhibitors of metabolism (24, 25), led us to hypothesise that these genes are operonic, a premise that we confirmed using co-transcriptional and growth dependent expression analyses. Our finding that PPE25 and PPE26 physically interact was not surprising given the commonality of interaction between the products of co-ordinately expressed genes. However, based on our experience working with PE-PPE clusters (16), we also expected to detect PPE25-PE18 and/or PPE26-PPE18 interactions in our assays, which we did not, highlighting the possible functional relevance of this specific interaction. This also represents the second identification of PPE-PPE interaction, providing credence to our earlier observation of interaction between PPE50 and PPE51 (26), a finding that had not been reported earlier. *M. smegmatis* strains expressing *PPE25, PPE26*, and *PE18* showed enhanced intra-macrophage survival compared to the control strain - with the increase in CFUs correlating with a decrease in *iNOS2* levels. Surprisingly, the recombinant strain expressing the entire operon did not show improved survival compared to controls, indicating a possible antagonistic role for the PPE25-PPE26 heterodimer in the independent CFU control function of the two proteins. *iNOS2* levels did not change over the control levels in this scenario, explaining the lack of CFU enhancement.

Cytokine measurements following infection of THP-1 macrophages with recombinant strains expressing the cluster genes individually or as a whole, showed that both PPE25 and PPE26 could mediate the intra-macrophage lowering of *IL-12* levels and an increase in *IL-10* expression in macrophages. Consistent with the established TLR2 binding activity of PPE26 (22), THP-1 macrophages infected with recombinant strains *M. smegmatis* expressing *PPE26 & PPE25-PE18-PPE26* showed enhanced p38 phosphorylation. It is highly likely that that the modulation of IL-10 and IL-12 by these proteins is mediated through the TLR2-p38-MAPK axis.

Our findings implicate the ESX-5 associated *PPE25-PE18-PPE26* gene cluster in *M. tb* pathogenesis *via* the modulation of pro- and anti-inflammatory cytokine expression, and by facilitating its intracellular survival. Evidence from this research also suggests that the PPE proteins possess unique functions in their monomeric form, and may have possible opposing functions when they heterodimerise with other PPE proteins (Fig. 6) – an observation we also made with the PPE50-PPE51 protein pair in the context of TLR1 binding (26). While the mechanism of this phenomenon remains to be elucidated, it represents a significant expansion in the immune-regulation arsenal of *M. tb*, and highlights the enormous versatility that this pathogen possesses in evading immune responses, to establish and persist in its human host.

**Figure 6:**
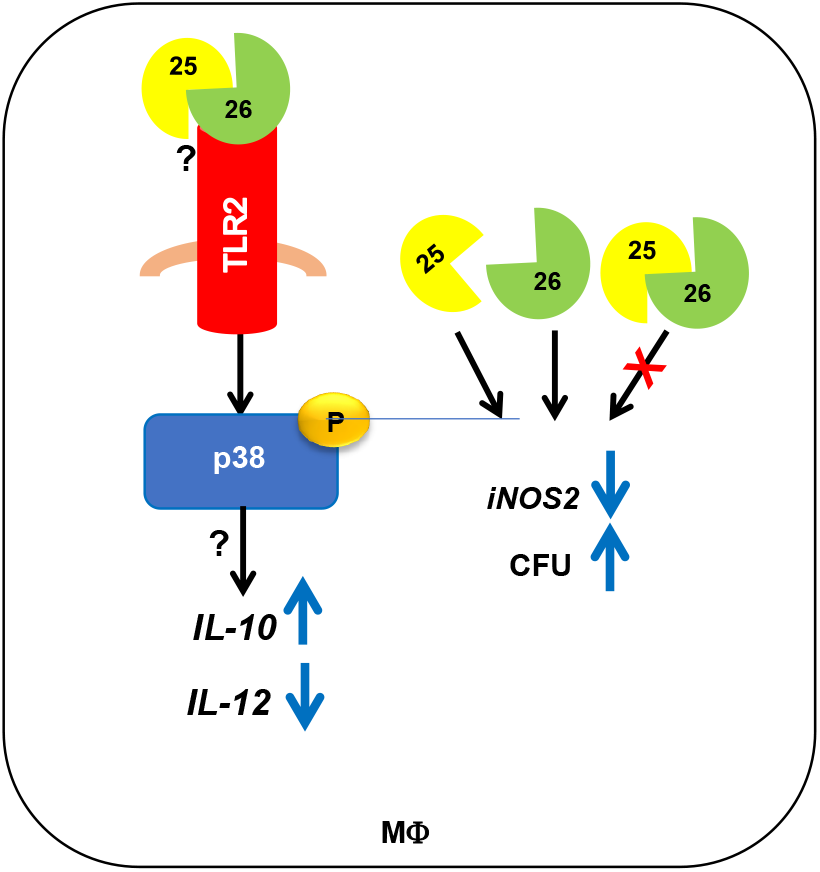
Summary of the study illustrating the immunomodulatory role of PPE25/PPE26. MΦ - Macrophage

## Author contributions

T. R. R. designed the study. R. P. M., K. P., and K. C. performed the experiments. T. R. R., R. P. M., K. P., and K. C. analysed the data. T. R. R. wrote the manuscript.

## References

1. Behar SM, Sassetti CM. Immunology: Fixing the odds against tuberculosis. Nature. 2014;511(7507):39–40.

2. DiGiuseppe Champion PA, Cox JS. Protein secretion systems in Mycobacteria. Cell Microbiol. 2007;9(6):1376–84.

3. Fishbein S, van Wyk N, Warren RM, Sampson SL. Phylogeny to function: PE/PPE protein evolution and impact on Mycobacterium tuberculosis pathogenicity. Mol Microbiol. 2015;96(5):901–16.

4. Abdallah AM, Gey van Pittius NC, DiGiuseppe Champion PA, Cox J, Luirink J, Vandenbroucke-Grauls CM, et al. Type VII secretion--mycobacteria show the way. Nat Rev Microbiol. 2007;5(11):883–91.

5. Bottai D, Di Luca M, Majlessi L, Frigui W, Simeone R, Sayes F, et al. Disruption of the ESX-5 system of Mycobacterium tuberculosis causes loss of PPE protein secretion, reduction of cell wall integrity and strong attenuation. Mol Microbiol. 2012;83(6):1195–209.

6. Sayes F, Sun L, Di Luca M, Simeone R, Degaiffier N, Fiette L, et al. Strong immunogenicity and cross-reactivity of Mycobacterium tuberculosis ESX-5 type VII secretion: encoded PE-PPE proteins predicts vaccine potential. Cell Host Microbe. 2012;11(4):352–63.

7. McNamara M, Danelishvili L, Bermudez LE. The Mycobacterium avium ESX-5 PPE protein, PPE25-MAV, interacts with an ESAT-6 family Protein, MAV_2921, and localizes to the bacterial surface. Microb Pathog. 2012;52(4):227–38.

8. Wu M, Li M, Yue Y, Xu W. DNA vaccine with discontinuous T-cell epitope insertions into HSP65 scaffold as a potential means to improve immunogenicity of multi-epitope Mycobacterium tuberculosis vaccine. Microbiol Immunol. 2016;60(9):634–45.

9. Majumdar SD, Vashist A, Dhingra S, Gupta R, Singh A, Challu VK, et al. Appropriate DevR (DosR)-mediated signaling determines transcriptional response, hypoxic viability and virulence of Mycobacterium tuberculosis. PLoS One. 2012;7(4):e35847.

10. Voskuil MI, Schnappinger D, Rutherford R, Liu Y, Schoolnik GK. Regulation of the Mycobacterium tuberculosis PE/PPE genes. Tuberculosis (Edinb). 2004;84(3-4):256-62.

11. Mi Y, Bao L, Gu D, Luo T, Sun C, Yang G. Mycobacterium tuberculosis PPE25 and PPE26 proteins expressed in Mycobacterium smegmatis modulate cytokine secretion in mouse macrophages and enhance mycobacterial survival. Res Microbiol. 2017;168(3):234–43.

12. Spratt JM, Ryan AA, Britton WJ, Triccas JA. Epitope-tagging vectors for the expression and detection of recombinant proteins in mycobacteria. Plasmid. 2005;53(3):269–73.

13. Cascioferro A, Delogu G, Colone M, Sali M, Stringaro A, Arancia G, et al. PE is a functional domain responsible for protein translocation and localization on mycobacterial cell wall. Mol Microbiol. 2007;66(6):1536–47.

14. Stover CK, de la Cruz VF, Fuerst TR, Burlein JE, Benson LA, Bennett LT, et al. New use of BCG for recombinant vaccines. Nature. 1991;351(6326):456–60.

15. Tiwari BM, Kannan N, Vemu L, Raghunand TR. The Mycobacterium tuberculosis PE proteins Rv0285 and Rv1386 modulate innate immunity and mediate bacillary survival in macrophages. PLoS One. 2012;7(12):e51686.

16. Tiwari B, Soory A, Raghunand TR. An immunomodulatory role for the Mycobacterium tuberculosis region of difference 1 locus proteins PE35 (Rv3872) and PPE68 (Rv3873). FEBS J. 2014;281(6):1556–70.

17. Singh A, Mai D, Kumar A, Steyn AJ. Dissecting virulence pathways of Mycobacterium tuberculosis through protein-protein association. Proc Natl Acad Sci U S A. 2006;103(30):11346–51.

18. Cserzo M, Wallin E, Simon I, von Heijne G, Elofsson A. Prediction of transmembrane alpha-helices in prokaryotic membrane proteins: the dense alignment surface method. Protein Eng. 1997;10(6):673–6.

19. Kyte J, Doolittle RF. A simple method for displaying the hydropathic character of a protein. J Mol Biol. 1982;157(1):105–32.

20. Akhter Y, Ehebauer MT, Mukhopadhyay S, and Hasnain SE. The PE/PPE multigene family codes for virulence factors and is a possible source of mycobacterial antigenic variation: perhaps more? Biochimie. 2012;94(1):110–6.

21. Flynn JL, Chan J. Immunology of tuberculosis. Annu Rev Immunol. 2001;19:93–129.

22. Su H, Kong C, Zhu L, Huang Q, Luo L, Wang H, et al. PPE26 induces TLR2-dependent activation of macrophages and drives Th1-type T-cell immunity by triggering the cross-talk of multiple pathways involved in the host response. Oncotarget. 2015;6(36):38517–37.

23. Ganguly N, Siddiqui I, Sharma P. Role of M. tuberculosis RD-1 region encoded secretory proteins in protective response and virulence. Tuberculosis (Edinb). 2008;88(6):510–7.

24. Boshoff HI, Myers TG, Copp BR, McNeil MR, Wilson MA, Barry CE, 3rd. The transcriptional responses of Mycobacterium tuberculosis to inhibitors of metabolism: novel insights into drug mechanisms of action. J Biol Chem. 2004;279(38):40174–84.

25. Mohareer K, Tundup S, Hasnain SE. Transcriptional regulation of Mycobacterium tuberculosis PE/PPE genes: a molecular switch to virulence? J Mol Microbiol Biotechnol. 2011;21(3-4):97-109.

26. Mukku RP, Poornima K, Yadav S, Raghunand TR. Delineating the functional role of the PPE50 (Rv3135) - PPE51 (Rv3136) gene cluster in the pathophysiology of Mycobacterium tuberculosis. Microbes Infect. 2024;26(3):105248.

