## Supplementary Data for "The *PPE25 (Rv1787) - PE18 (Rv1788) - PPE26 (Rv1789)* gene cluster encodes an interacting protein pair and is involved in immune evasion by *Mycobacterium tuberculosis*"

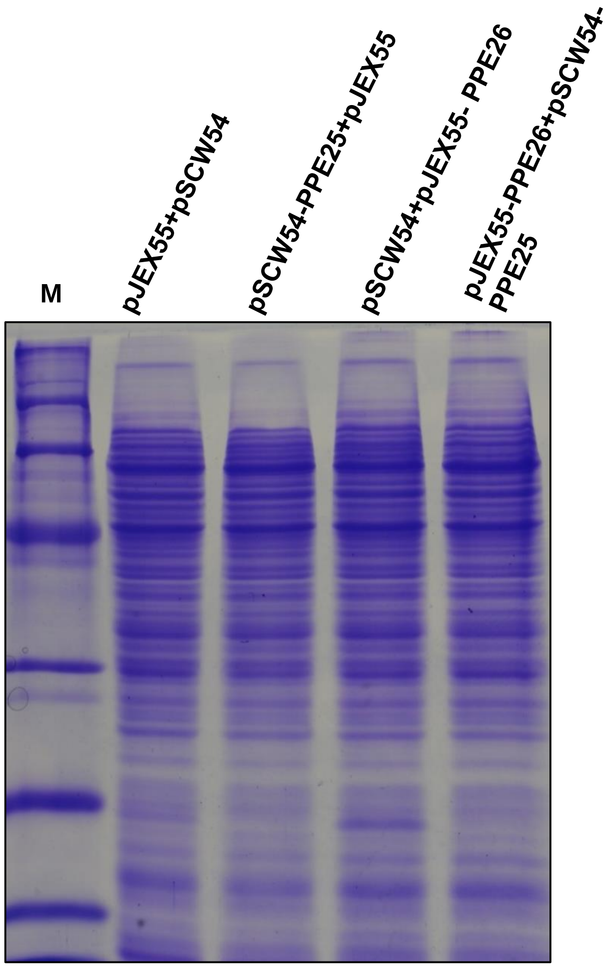


**Figure S1**: ***In vivo* co-IP of PPE25-PPE26.** SDS-PAGE profiles showing equal loading for *in vivo* co-IP reactions of *M. smegmatis* expressing PPE25-6xHis and PPE26-c-myc, precipitated with an anti-c-myc antibody and probed with an anti-His antibody.

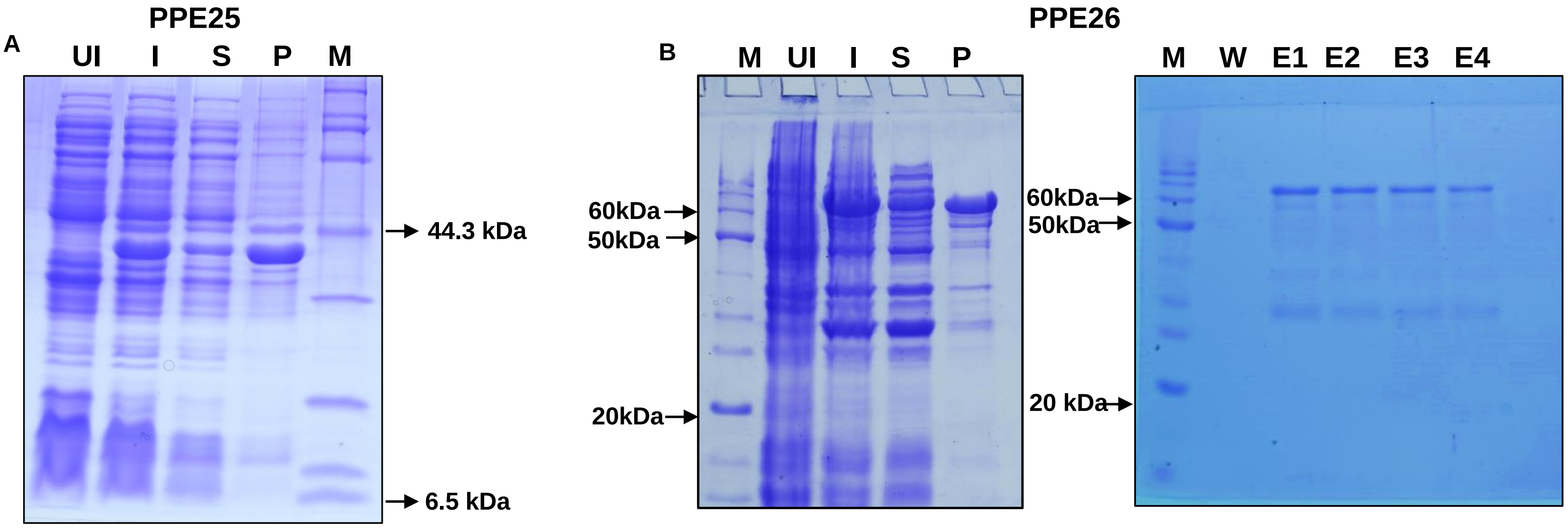


**Figure S2: Expression profile of PPE25 and PPE26. (**A) Expression profile of PPE25 in *E. coli* BL21DE3. (B) Expression (left panel), and GSH based affinity chromatography purification profile (right panel) of PPE26 expressed in *E. coli* BL21DE3. UI - Uninduced sample, I - Induced sample, S - Supernatant, P - Pellet Fraction, M - Marker, W - Wash, E - Eluate.

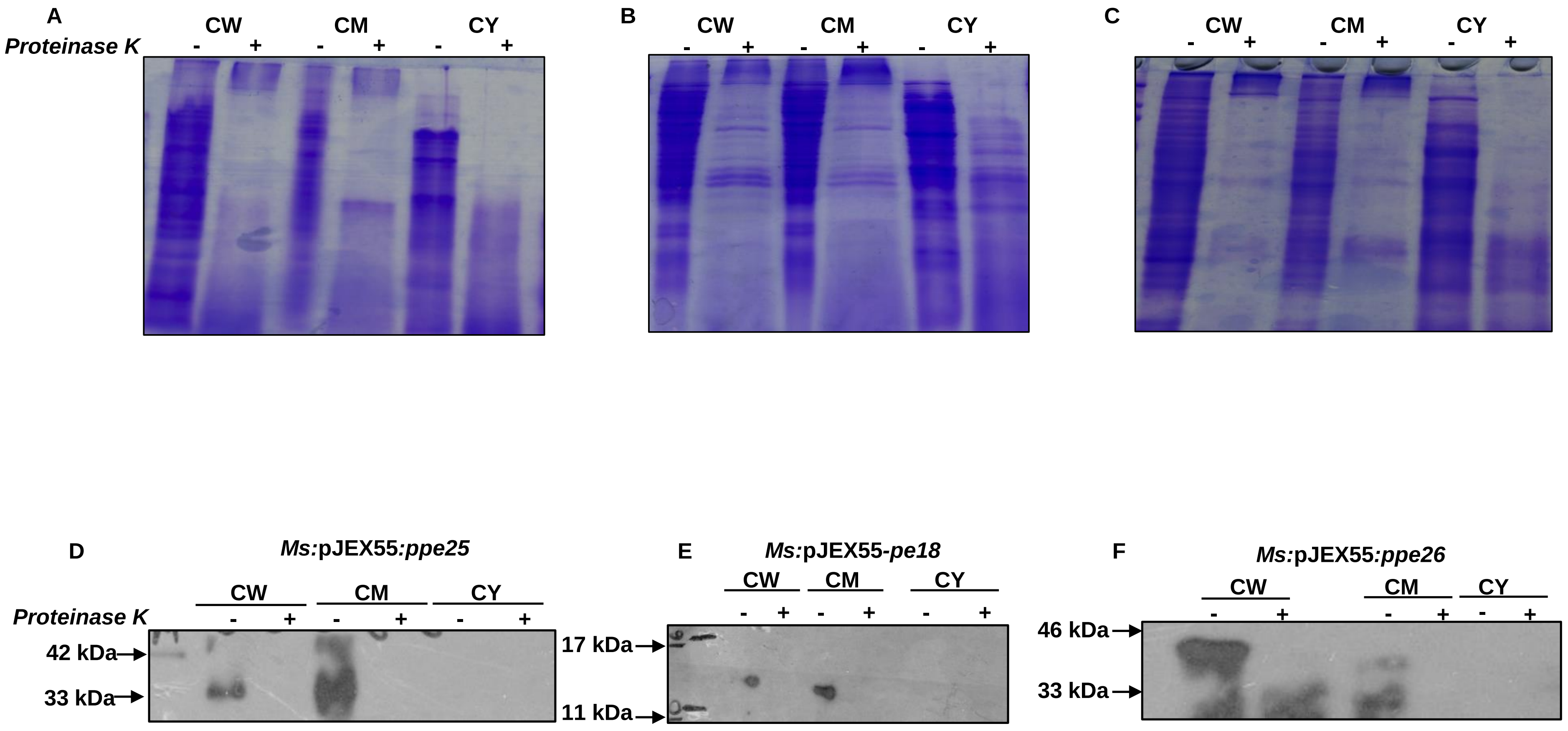


**Figure S3 Subcellular localisation and surface accessibility.** Coomassie blue stained SDS-PAGE profiles of Proteinase K treated *M. smegmatis* expressing PPE25-c-myc (A), PE18-c-myc (B), and PPE26-c-myc (C), demonstrating equivalent loading of the sub-cellular fractions. Immunodetection of PPE25 (D), PE18 (E), and PPE26 (F), in subcellular fractions (CM - Cell Membrane, CW - Cell Wall, CY - Cytoplasm). An anti-c-myc monoclonal antibody was used to identify all proteins.

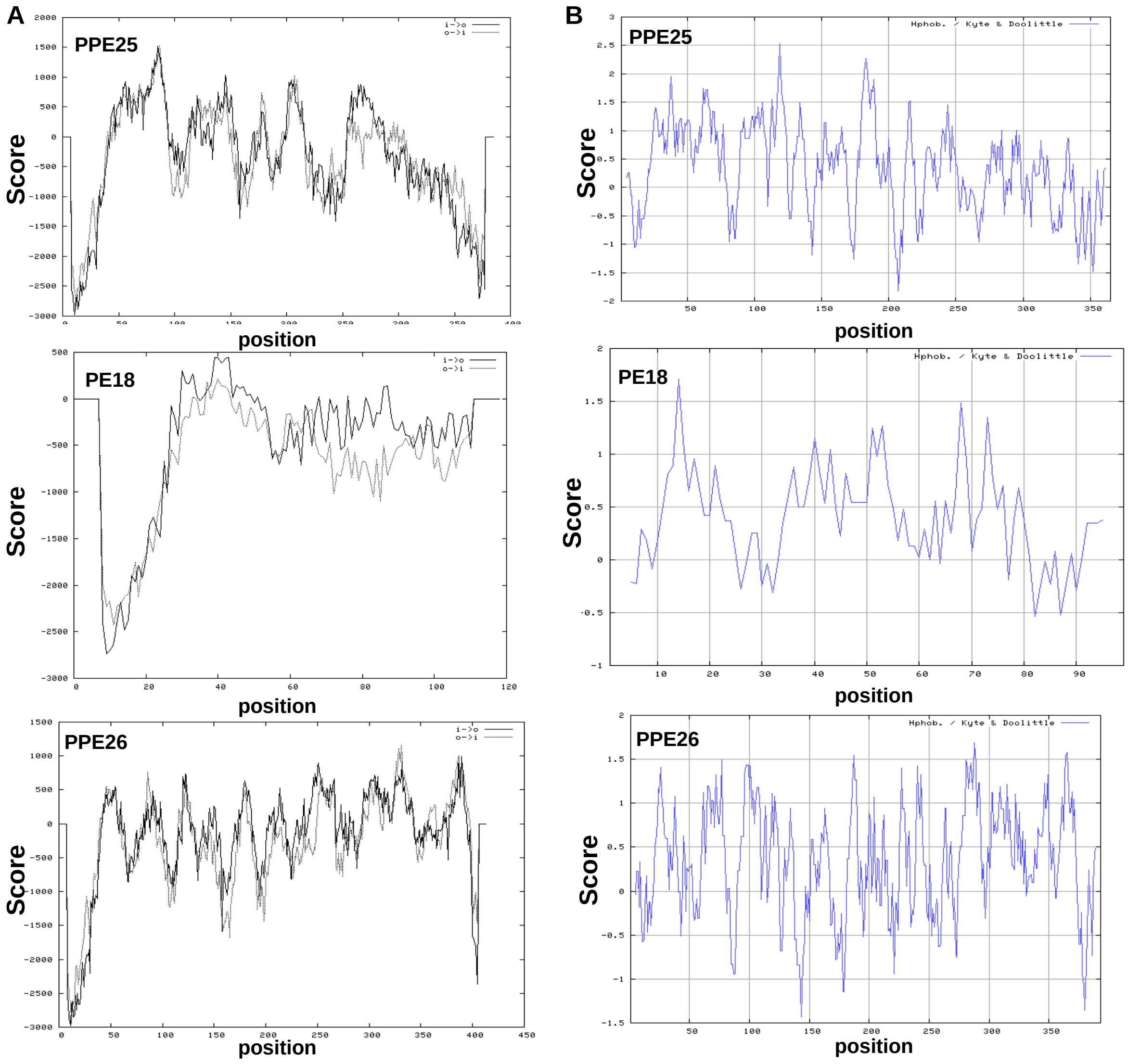


**Figure S4: *in silico* protein sequence analysis of *M. tb* PPE25, PPE18, and PPE26.**
Analyses of PPE25, PPE18, and PPE26 protein sequences for transmembrane prediction (A) and hydrophobicity (B).

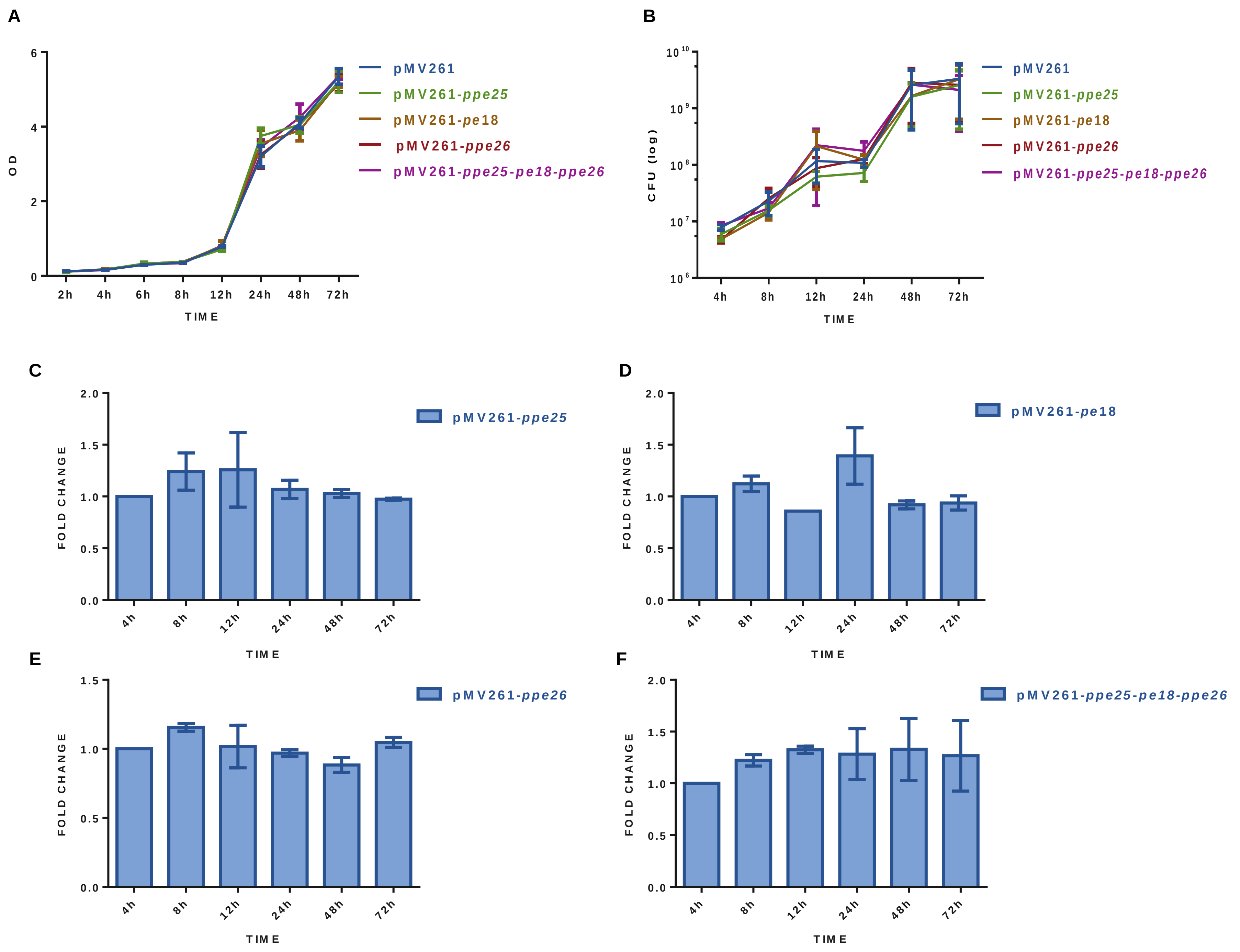


**Figure S5: Gene expression and growth analysis of recombinant *Mycobacterium smegmatis* strains expressing *PPE25*, *PPE18*, and *PPE26*.** *In vitro* growth profiles of *M. smegmatis* expressing pMV261, *PPE25*, *PE18*, *PPE26*, and *PPE25-PE18-PPE26* (A) OD, (B) CFU counts. Transcript levels of *PPE25* (C), *PPE18* (D), *PPE26* (E), and *PPE25-PE18-PPE26* (F), were measured by real-time RT-PCR as a function of growth. Data are shown as the mean ± SEM of transcript levels normalised to the mRNA level of the relevant gene at time 4h (value = 1).

**Table S1: Primers used in this study**

| | | |
|---|---|---|
| pMV261-PPE25F | FP for cloning *PPE25* in pMV261 | 5’CGCGAATTCTTGGACTTCGGGGCGTTACC3’ |
| pMV261-PPE25R | RP for cloning *PPE25* in pMV261 | 5’CCCAAGCTTTTATCCGGCCGATGGCGGG3’ |
| pMV261-PE18F | FP for cloning *PE18* in pMV261 | 5’CCGGAATTCCATGTCGTTTGTGACTACCCAACCAGAAG3’ |
| pMV261-PE18R | RP for cloning *PE18* in pMV261 | 5’CCCAAGCTTCTAGCCGGCCGCGGCCGC3’ |
| pMV261-PPE26F | FP for cloning *PPE26* in pMV261 | 5’CGCGGATCCATGGATTTTGGGGCGTTGCCG3’ |
| pMV261-PPE26R | RP for cloning *PPE26* in pMV261 | 5’CCCAAGCTTCTATCCGGCGAAGGGTGGG3’ |
| pJEX55-PPE25F | FP for cloning *PPE25* in pJEX55 | 5’CGCGGATCCTTGGACTTCGGGGCGTTACC3’ |
| pJEX55-PPE25R | RP for cloning *PPE25* in pJEX55 | 5’CCCGAATTCTTATCCGGCCGATGGCGGG3’ |
| pJEX55-PPE26F | FP for cloning *PPE26* in pJEX55 | 5’CGCGGATCCATGGATTTTGGGGCGTTGCCG3’ |
| pJEX55-PPE26R | RP for cloning *PPE26* in pJEX55 | 5’CCCGAATTCCTATCCGGCGAAGGGTGGG3’ |
| pJEX55-PE18F | FP for cloning *PE18* in pJEX55 | 5’CCGGGATCCCATGTCGTTTGTGACTACCCAACCAGAAG3’ |
| pJEX55-PE18R | RP for cloning *PE18* in pJEX55 | 5’CCCGAATTCCTAGCCGGCCGCGGCCGC3’ |
| pET22b-PPE25-F | FP for cloning *PPE25* in pET22b | 5’GGGAATTCCATATGTTGGACTTCGGGGCGTTACC3’ |
| pET22b-PPE25-R | RP for cloning *PPE25* in pET22b | 5’CCGCTCGAGCGGCTCGAGAGGTGGAGTGCCAGCGGTGT3’ |
| pGEX-6P1-PPE26-F | FP for cloning *PPE26* in pGEX-6p1 | 5’CGCGGATCCATGGATTTTGGGGCGTTGCCG3’ |
| pGEX6p1-PPE26-R | RP for cloning *PPE26* in pGEX-6p1 | 5’CCGCTCGAGTCAGTGGTGGTGGTGGTGGTGTCCGGCGAAGGGTGGGCG3’ |
| pUAB300-PPE25F | FP for MPFC analysis of PPE25-PPE26 | 5’CGCGAATTCTTGGACTTCGGGGCGTTACC3’ |
| pUAB300-PPE25R | RP for MPFC analysis of PPE25-PPE26 | 5’CCCAAGCTTTTATCCGGCCGATGGCGGG3’ |
| pUAB400-PPE25F | FP for MPFC analysis of PPE25-PPE26 | 5’CCGGAATTCCTTGGACTTCGGGGCGTTACC3’ |
| pUAB400-PPE25R | RP for MPFC analysis of PPE25-PPE26 | 5’CCCAAGCTTTTATCCGGCCGATGGCGGGC3’ |
| pUAB400-PE18F | FP for MPFC analysis of PPE25-PE18 &PPE26-PE18 | 5’CCGGAATTCCATGTCGTTTGTGACTACCCAACCAGAAG3’ |

| | | |
|---|---|---|
| pUAB400-PE18R | RP for MPFC analysis of PPE25-PE18 &PPE26-PE18 | 5’CCCAAGCTTCTAGCCGGCCGCGGCCGC3’ |
| pUAB300-PPE25F | FP for MPFC analysis of PPE25-PE18 | 5’CGCGAATTCTTGGACTTCGGGGCGTTACC3’ |
| pUAB300-PPE25R | RP for MPFC analysis of PPE25-PE18 | 5’CCCAAGCTTTTATCCGGCCGATGGCGGG3’ |
| pUAB300-PPE26F | FP for MPFC analysis of PPE26-PE18 & PPE25-PPE26 | 5’CGCGGATCCATGGATTTTGGGGCGTTGCCG3’ |
| pUAB300-PPE26R | RP for MPFC analysis of PPE26-PE18 & PPE25-PPE26 | 5’CCCAAGCTTCTATCCGGCGAAGGGTGGG3’ |
| pUAB400-PPE25F | FP for MPFC analysis of PPE25-PPE26 | 5’CCGGAATTCCTTGGACTTCGGGGCGTTACC3’ |
| pUAB400-PPE25R | RP for MPFC analysis of PPE25-PPE26 | 5’CCCAAGCTTTTATCCGGCCGATGGCGGGC3’ |
| RT-PPE25F | FP for expression analysis of *PPE25* in *M. smegmatis* (RT PCR) and co-operonic analysis | 5’TTGGACTTCGGGGCGTTAC3’ |
| RT-PPE25R | RP for expression analysis of *PPE25* in *M. smegmatis* (RT PCR) and co-operonic analysis | 5’GCCACCATCGACAACGAC3’ |
| junction1RTF | FP Co-operonic analysis for PPE25-PE18-PPE26 and co-operonic analysis | 5’ATATGGGTTCCGTCACAGTGT3’ |
| junction1RTR | RP Co-operonic analysis for PPE25-PE18-PPE26 and co-operonic analysis | 5’ATGCGGAGCCGATTCCCT3’ |
| RT-PE18F | FP for expression analysis of *PE18* in *M. smegmatis* (RT PCR) and co-operonic analysis | 5’ATGTCGTTTGTGACTACCCAAC3’ |
| RT-PE18R | RP for expression analysis of *PE18* in *M. smegmatis* (RT PCR) co-operonic analysis | 5’ATGTCGTTTGTGACTACCCAAC3’ |
| junction2RTF | FP Co-operonic analysis for PPE25-PE18-PPE26 | 5’GTCGTATGCTGCTACCGAG3’ |
| junction2RTR | RP Co-operonic analysis for PPE25-PE18-PPE26 | 5’GTCTCATAACCGGTGGCC3’ |
| RT-PPE26F | FP for expression analysis of *PPE26* in *M. smegmatis* (RT PCR) and co-operonic analysis | 5’ATGGATTTTGGGGCGTTGCCG3’ |

| | | |
|---|---|---|
| RT-PPE26R | RP for expression analysis of *PE18* in *M. smegmatis* (RT PCR) co-operonic analysis | 5’ATCGCCGCTGACGCCGGA 3’ |
| huIL10 RT –F | FP for RT PCR of human IL-10 | 5'CCTTGTCTGAGATGATCCA GTT3' |
| huIL10 RT –R | RP for RT PCR of human IL-10 | 5'TAAAGGCATTCTTCACCTG CTC3' |
| huActin β RT-F | FP for RT PCR of human β-actin | 5'GAGCAAGAGAGGCATCCT CAC3' |
| huActin β RT-R | RP for RT PCR of human β-actin | 5'CTCAAACATGATCTGGGTC ATC3' |
| huIL12b RT-F | FP for RT PCR of human IL-12b | 5'ATCAGGGACATCATCAAAC CTG3' |
| huIL12b RT-R | RP for RT PCR of human IL-12b | 5'AGGTCTTGTCCGTGAAGAC TC3' |
| hu RT INOS-F | FP for RT PCR of human *iNOS2* | 5’AGTTTCCAGAAGCAGAATG TGAC3’ |
| hu RT INOS-R | RP for RT PCR of human *iNOS2* | 5’GTAGAAAGGGGACAGGAC GTA3’ |
| *MssigA* RT-F | FP for RT PCR of *M. smegmatis sigA* | 5’GCCAGCTCGGTGACTTCA3 ’ |
| *MssigA* RT-R | RP for RT PCR of *M. smegmatis sigA* | 5’CGTGACGCCGTAGACCTG3 ’ |
